# Time to Potential Collision: A Dynamic Approach To Study Vessel-Whale Close Encounters

**DOI:** 10.64898/2026.03.23.713354

**Authors:** Rita Santos, Cláudia Oliveira-Rodrigues, Iolanda M. Silva, Raul Valente, Luís Afonso, Ágatha Gil, Catarina Vinagre, Annalisa Sambolino, Marc Fernández, Filipe Alves, Isabel Sousa-Pinto, Ana Mafalda Correia

## Abstract

Vessel-whale collisions are a growing global concern and remain challenging to quantify. Therefore, the use of proxies, such as Close Encounters (CEs) that comprise Surprise Encounters (SEs) and Near-Miss Events (NMEs), has been proposed and widely employed to assess collision risk. To better understand this risk in the Eastern North Atlantic, where maritime traffic is intensive, this study aimed to redefine and quantify CEs, and to assess detectability-related variables that may affect CE identification. CEs were assessed using a cetacean occurrence dataset collected between 2012 and 2024 on board cargo ships and oceanographic vessels. CEs thresholds were redefined based on Time to Potential Collision (TPC), rather than distance alone (as described in literature), to allow a more dynamic, risk-based, and speed-sensitive approach. In total, 1226 sightings of whales (baleen, sperm, and beaked whales) were recorded, of which 37.4% were classified as SEs and 2.0% as NMEs. The sperm whale, *Physeter macrocephalus*, was the species most frequently involved in CEs (13.9% of all CEs), followed by the Cuvier’s beaked whale, Zip*hius cavirostris* (11.8%). A Generalized Additive Model was used to assess the influence of detectability-related variables (i.e., meteorological conditions, whale taxa, vessel characteristics, and Marine Mammals Observers (MMOs) experience) on TPC. Significantly lower TPC values were observed with beaked whales, cargo ships, poor visibility conditions, and less experienced MMOs. The results of this study provide an CE’s assessment in this region and contribute to the ongoing efforts to standardize CE quantification, by using TPC as a metric. This work also highlights the importance of decreased speeds and the presence of experienced MMOs on board to increase detection probability and TPC, thereby potentially minimizing collision risk.

## Introduction

Over recent decades, marine megafauna have been exposed to numerous anthropogenic threats, with ship strikes becoming a growing global concern (Laist et al., 2001; Van Waerebeek et al., 2007; Ritter, 2012; Ritter et al., 2016; Currie et al., 2017; Simmonds, 2018; Schoeman et al., 2020; Nisi et al., 2024). Particularly, vessel-whale collisions are a major threat to cetaceans, especially for large whales (Laist et al., 2001; Cates et al., 2017; IWC,2022; Vighi, 2025). Collisions occur worldwide, with a higher risk in areas where core cetacean habitats overlap with intensive vessel traffic, a situation that has increased over time (Cates et al., 2017; Schoeman et al., 2020; Winkler et al., 2020; Nisi et al., 2024). Collision risk is intricately linked to vessel characteristics, namely vessel size and speed. Most serious and lethal strikes occur in small vessels (<20 m) travelling above 14 knots, while in large vessels (>20 m) collisions are likely to be fatal regardless of speed (Laist et al., 2001; Vanderlaan & Taggart, 2007; Conn & Silber, 2013; Kelley et al., 2020; Blondin et al., 2025; Garrison et al., 2025). Additionally, all species can potentially be hit, but lone, sick whales, and individuals engaged in feeding, resting, or socializing activities, may be oblivious to boat presence, placing them at a higher risk of collision (Laist et al., 2001; Dolman et al., 2006; Ritter et al., 2016; Currie et al., 2017; Blondin et al., 2025).

Assessing collision likelihood is crucial for identifying high-risk areas and implementing suitable mitigation strategies (Schoeman et al., 2020). Despite some economic concerns, speed limits are considered effective measures to decrease the likelihood of a strike, by allowing more time for evasive maneuvers upon whale detection, or reduce its lethality (Vanderlaan and Taggart, 2007; Gende et al., 2011; Conn & Silber, 2013; Laist et al., 2014; Redfern et al., 2024). Nonetheless, recent studies suggest that speed regulations for large vessels are less effective, or even ineffective, in contrast to those applied in small vessels (Kelley et al., 2020; Blondin et al., 2025; Garrison et al., 2025). Studies also suggest that the first approach should be minimizing the vessel-whale overlap, by re-routing shipping lanes or defining areas to avoid (Vanderlaan et al., 2008; IWC, 2011; IMO, 2016; Frantzis et al., 2019). Moreover, the employment of dedicated Marine Mammal Observers (MMOs) is considered an efficient way to prevent collisions, particularly when whales and vessels inevitably overlap (Weinrich et al., 2010; Wiley et al., 2016; Flynn & Calambokidis, 2019). By detecting the animals beforehand, there is more time to take on precautionary measures potentially needed to avoid a strike (Carrillo & Ritter, 2010; Weinrich et al., 2010; IWC, 2011; Ritter & Panigada, 2019). MMOs also contribute to the on-going effort to collect data on collisions, allowing a better understanding of these incidents (Weinrich et al., 2010; Gende et al., 2011; IWC, 2011).

To fully understand the scale of this problem, there is a need to accurately quantify ship strikes. However, the lack of data regarding the location and magnitude of this issue poses a barrier to the assessment of the problem (Dolman et al., 2006; IWC, 2011; Ritter & Panigada, 2019). On large vessels, especially cargo ships, visibility towards the bow is frequently limited, as the bridge is usually positioned towards the stern, and, as a result, the crew is often unaware of cetaceans’ presence in the vessel’s path, which makes many collisions go unnoticed (Laist et al., 2001; IWC, 2022; Vighi, 2025). A lack of awareness of the problem and how to correctly report it also contributes to underreporting (Laist et al., 2001; Dolman et al., 2006; Van Waerebeek et al., 2007; Van Waerebeek & Leaper, 2008; Ritter & Panigada, 2019). The reluctance to report collisions (or suspected collisions), often driven by fear of reprisals, fines or damage to reputation, is an additional contributor to the underestimation of vessel strikes (Neilson et al., 2012; Ransome et al., 2021). Moreover, despite direct observations being valuable to collect relevant data, *in situ* records are scarce, and a large amount of data on ship strikes comes from examinations of stranded or recovered carcasses (IWC, 2011; Winkler et al., 2020; Vighi, 2025). This represents only a small fraction, as many carcasses sink, and non-lethal strikes are not being considered (Dolman et al., 2006; IWC, 2011).

As a result, ship strike numbers are very likely underestimated, and the frequency of vessel-whale collisions appears to be increasing, although it remains unclear whether this reflects a true rise or simply improved reporting, making the extent of this issue extremely challenging to evaluate (Ritter & Panigada, 2019; Winkler et al., 2020; Ransome et al., 2021). Given this uncertainty, the use of indexes such as Surprise Encounters (SEs) and Near-Miss Events (NMEs), has been proposed to assess collision risk (Richardson et al., 2011; Ritter et al., 2016; David et al., 2022). However, there are different approaches and definitions when considering these events. The definition of a SE was initially proposed by Richardson et al. (2011), corresponding to an unexpected close encounter between a vessel and a whale at a distance below 300 m from the vessel, while an NME corresponded to a sighting occurring near the vessel’s bow at a distance below 80 m. The term NME has been more widely used and generally defined as an unplanned close encounter that did not involve any confirmed collision but could potentially have resulted in one, if whale or vessel had not taken action to avoid it (Ritter et al., 2016; Stack et al., 2016). A clear definition is still required, as these collision risk indexes are not being recorded in a standard way, if they are recorded at all (IWC, 2011; Ritter et al., 2016; Stack et al., 2016). Without a precise characterization, quantifying SEs and NMEs and comparing studies can become particularly challenging and subjective (Ritter et al., 2016). This lack of consistency underscores the urgent need for the development of consistent definitions and monitoring protocols to improve our understanding and mitigation of vessel-whale collisions.

Therefore, the aim of the present study is to redefine the basis of SEs and NMEs, which have been previously defined based on vessel-whale distance, and to propose an approach for the standardized recording of collision risk proxies, accounting for the vessel speed, based on Time to Potential Collision (TPC). To that, we quantify these events in the Eastern North Atlantic (ENA) as a case study, based on a longitudinal dataset. As specific aims, we investigate how different factors, such as meteorological conditions, taxonomic family, vessel characteristics, and experience of MMOs, influence the TPC of SEs and NMEs. By enhancing our understanding of vessel-whale risk interactions in the ENA, providing a method for recording them worldwide, and assessing the key variables that may influence detection, this study will provide valuable insights to support the development of management and conservation strategies aimed at reducing collision risk.

## Methods

### Study Area

The present study was conducted in the ENA (Fig. 1), encompassing the waters off the Iberian Peninsula, the Macaronesia Archipelagos, and Northwest Africa. Four main routes were monitored on board cargo ships: mainland Portugal – Azores, mainland Portugal – Madeira, mainland Portugal – Azores – Madeira and mainland Portugal – Cape Verde, with stopovers in the Canary Islands, Senegal, Mauritania, and Northwest Spain. Cetacean monitoring on board oceanographic vessels was conducted off mainland Portugal and over the Azores and Madeira archipelagos. The study area is characterized by a complex and dynamic oceanography, along with a varied topography (Mason, 2009; Mártinez, 2021). The overall diverse seafloor and depth of the region contribute to high cetacean diversity and shapes their distribution (Waring *et al*., 2009; Correia et al., 2019, 2020; Cartagena-Matos et al., 2021). Cetaceans are a vital component of the study area and are widespread throughout the ENA (Correia et al., 2020; Cartagena-Matos et al., 2021).

**Figure 1.**
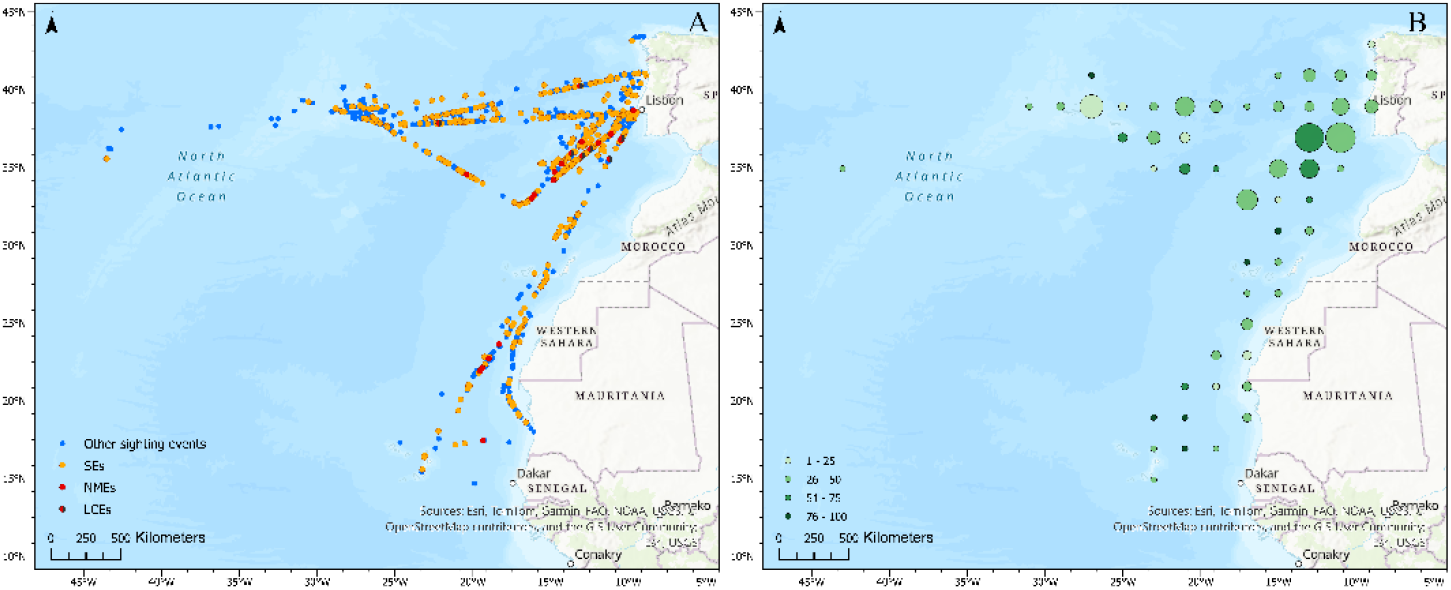
Study area with: (A) spatial distribution of Surprise Encounters (SEs), Near-Miss Events (NMEs), Likely Collision Events (LCEs), and other whale sighting events; and (B) proportion of Close Encounters (CEs) in relation to the total number of sightings per grid cell. Circle size represents the total count of sightings per grid cell, and the circle’s color intensity reflects the proportion of CEs relative to the number of sightings within each cell. Data were collected during cetacean monitoring surveys conducted between 2012 and 2024 using cargo ships and oceanographic vessels as research platforms.

The ENA is a very busy shipping region and, between 1820 and 2019, most of the reported ship strikes occurred in the North Atlantic Ocean (Winkler et al., 2020; Robbins et al., 2022). Specifically, the Canary Islands are one of the most affected regions worldwide, where collisions seem to pose a serious threat to cetaceans (Winkler et al., 2020; Carrillo & Ritter, 2010). Moreover, the Azores and Madeira archipelagos are positioned close to several marine traffic corridors and routes (used by cruise and cargo ships, and leisure boats), with many different destinations (Cunha et al., 2017; Soares et al., 2020). Similarly, the coast of mainland Portugal is crossed by numerous fishing vessels, cargo and cruise ships, and leisure crafts, creating a high concentration of maritime traffic (Silveira et al., 2012, 2013).

### Data collection

Data were collected through visual monitoring conducted on board four cargo ships – Insular (IN), Lagoa (LA), Monte Brasil (MB), and Monte da Guia (MG) – and two oceanographic vessels – NRP Almirante Gago Coutinho (GC) and NRP D. Carlos I (DC). Surveys were conducted between 2012 and 2024, with a temporary interruption in cargo ships’ embarkations from 2020 to 2022 due to the COVID-19 pandemic. Monitoring primarily took place during summer and early autumn (July to October). Data were collected by two dedicated MMOs positioned on the wings of the navigation bridge, from sunrise to sunset, and following a standardized protocol for line-transect sampling (Correia et al., 2019). The recording apps (“MyTracks”, https://my-tracks.pt.aptoide.com/; “LocusMap”, https://www.locusmap.app/; and “ILogWhale”, EcoStrim, Interreg Maritime, CIMA Research Foundation) automatically recorded vessel speed associated with the location points, both manually and automatically added throughout the route. Upon cetacean sighting, key data were recorded, including the distance to the animal/group (i.e., measured in reticles below the horizon on the binocular’s scale), sighting angle, taxa, best estimate of group size, and the animal’s response to the vessel (indifferent, approach or avoiding). In addition to cetacean occurrence data, meteorological conditions were assessed by evaluating sea state (according to the Douglas scale), wind state (according to the Beaufort scale), and visibility (according to a categorical scale). Visibility was classified as low (<1 km), medium (1 – 4 km), good (4 – 10 km), or optimum (>10 km). On-effort (i.e., active dedicated monitoring by MMOs) transects were suspended whenever conditions became unfavorable for cetacean monitoring (e.g., Beaufort or Douglas values over 4, or low visibility), or when it was not possible to remain on the navigation bridge (e.g., instructions from the vessel’s commander, safety drills, or deck cleaning). Accordingly, cetacean sightings recorded during off-effort periods were considered opportunistic (Correia et al., 2019).

### Data processing

Initially, data from each survey were processed separately and were subsequently organized, structured, and cleaned (e.g., conversion of variables, insertion of codes, manual correction of any detected errors) to ensure consistency and accuracy. The sighting dataset was truncated to only include baleen and medium-to-large toothed whales, namely the sperm whale (Physeteridae), pygmy and dwarf sperm whales (Kogiidae), and beaked whales (Ziphiidae). For purposes of simplification, throughout this study, included species will be referred to as “whales”. Delphinidae species were excluded from the analysis as they are known for more structured group behaviors, are more active at the surface, tend to approach vessels, and are arguably less impacted by vessel collisions (Tyack, 1986; Van Waerebeek & Leaper, 2008; Hawkins & Gartside, 2009; Würsig, 2009; Jefferson & LeDuc, 2018; Winkler et al., 2020). Furthermore, whenever possible, sighting validations (i.e., species identification) were subsequently made through photographic records (Oliveira-Rodrigues et al., 2022). This was the case for all *Mesoplodon* genus sightings that include species difficult to distinguish at sea (e.g., *M. bidens* and *M. europaeus*).

To assess distance to the sightings, the Radial Distance (m), defined as the maximum distance from the observer to the animal, was estimated using the following formula (Heinemann, 1981; Lerczak & Hobbs, 1998; Marine Mammal Observer Association, 2024):

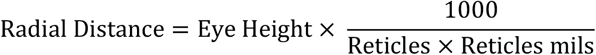

Eye Height refers to the sum of the observation deck’s height (m) and the average human eye level (1.59 m; First in Architecture, 2019). Reticles correspond to the number of binocular reticles measured from the horizon to the sighting, and Reticle mils are provided under the binocular’s user manuals. Using the Radial Distance, the distance between the animal and the vessel, hereafter Vessel-to-Whale Distance (m), was calculated. For that purpose, sightings were first categorized based on their relative position: either at the vessel’s bow or abeam. To do so, the angular range corresponding to the bow area (α) was calculated (Fig. S1), using each vessel’s dimensions, by using the following formula:

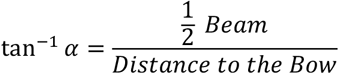

Distance to the bow refers to the distance between the observer and the bow of the vessel and Beam corresponds to the width of the vessel at its widest point. Based on this calculation, the bow area was defined as the angular range between -6º and 6º from the vessel’s heading for cargo ships, and between -23º and 23º for oceanographic vessels. The measures were determined to be the widest angle, rounded to the highest unit. The Vessel-to-Whale Distance was then calculated using the Pythagorean theorem. For sightings observed at the bow (Fig. S2A), the formula below was applied:

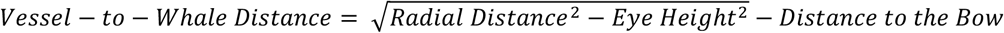

For the sightings occurring abeam (Fig. S2B), the Vessel-to-Whale Distance was estimated using the following formula:

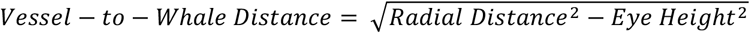

### Defining Close Encounters

In the context of this study, and throughout the remaining text, SEs and NMEs will be collectively referred to as Close Encounters (CEs). To define CEs, only sightings occurring between -90º and 90º relative to the vessel’s heading were considered. We used the approach described by Richardson et al. (2011), and later adopted in other studies (e.g., Stack et al., 2013; Currie et al., 2014; Currie et al., 2017), as a baseline for this study’s definition. According to Richardson et al. (2011) framework, CEs were defined based solely on the distance to the whale: a SE occurs when the distance is less than 300 m, while an NME occurs when the whale is located at the vessel’s bow and the distance is less than 80 m. In the present study, as vessel speed varied considerably (ranging from 1 to 22 knots, approximately), thresholds were defined based on Time to Potential Collision (TPC) rather than distance alone. TPC (s) essentially corresponds to the available time for the vessel to undertake evasive maneuvers (Stack et al., 2016), and it was estimated by dividing the Vessel-to-Whale Distance by the vessel’s speed (m/s), at the time of the sighting. When speed data were unavailable at the time of the sighting (i.e., missing data due to the app failing to automatically retrieve information, which occurred in 60 sighting events), the average speed (calculated with data from all available speed values during whale sightings) of that vessel was used, as ships usually maintain consistent cruise speeds. Given that Stack et al. (2013) reported a similar range of vessel speeds, the TPC corresponding to their lowest vessel speed (5 knots) for 300 meters and 80 meters was used to define the maximum TPC (threshold) for this study’s definition of SEs and NMEs, respectively. This conservative approach ensures that all potential CEs were included and is in line with Richardson et al. (2011), who provided the distance thresholds, but also consistent with Stack et al. (2016), who highlighted the speed range of vessels and emphasized the importance of considering maneuvering time to assess collision risk. Accounting for the conditions mentioned above, within the scope of this study, CEs were defined as whale sightings occurring near the vessel, between -90º and 90º relative to the vessel’s heading, and were categorized as follows: SEs, defined as the sightings with a TPC > 31.1 and ≤ 116.6s; and NMEs, defined as sightings with a TPC of ≤ 31.1s. Moreover, a sub-category for the NMEs occurring at the vessels’ bow area was also proposed as being Likely Collision Events (LCEs). LCEs represent the highest risk scenario as whales emerge right at the vessel’s bow, leaving limited time for detection and maneuvering.

### Descriptive and Spatial Analysis

Descriptive statistics were used to explore the general characteristics of the dataset in RStudio (R Version 4.3.2). For spatial analysis, ArcGIS Pro 3.4.3 was used to map CEs and other sighting events, and to calculate the proportion of CEs in relation to the number of sightings. The coordinate system used for all maps was World Mercator, projected from WGS84 coordinates (EPSG:4326).

### Modeling

A Generalized Additive Model (GAM) was fitted in RStudio (R Version 4.3.2) to evaluate the influence of detectability-related variables on TPC. Collated sightings included those registered during on-effort and off-effort periods. Firstly, a Mann-Whitney-Wilcoxon test was conducted to investigate any statistically significant differences between the TPC of sightings on-effort and off-effort to evaluate whether all sightings should be used in the modelling process (i.e., no statistically significant differences observed), or if off-effort sightings should be excluded from modelling (i.e., statistically significant differences observed). The level of significance was set at 0.05. The response variable of the model was the sightings’ TPC and being a continuous positive factor with a non-normal distribution, a gamma error distribution with a log link function was used (Dunn & Smyth, 2018; Pedersen et al., 2019). Detectability-related explanatory variables included sea and wind states, visibility, taxonomic family, group size, vessel (IN, LA, MB, MG, GC, or DC), vessel type (cargo ship or oceanographic vessel), vessel’s height, mean vessel speed, vessel’s beam, vessel’s distance from the observer to the bow, evaluation of the Least Experienced Observer (LEO) on board, evaluation of Most Experienced Observer (MEO) on board, average evaluation of the MMOs on board, and cumulative evaluation of the MMOs on board. LEO and MEO are performance metrics used to represent observer experience. MMOs experience was evaluated using qualitative criteria and assigned a score from 0 to 20, following Oliveira-Rodrigues et al. (2022). Prior to modeling, categories with very low sample sizes (<10) were removed from categorical variables to prevent highly unbalanced classes (e.g., sea and wind states equal to 0, Kogiidae sightings, and sightings on board GC). We used the Generalized Variance Inflation Factor (GVIF) to assess multicollinearity among predictors, which is appropriate for variables with multiple degrees of freedom (categorical predictors), considering a threshold of 5 (Hendrickx et al., 2004; Shikatani & Richman, 2024). For the GVIF assessment, vessel-related and MMO-related variables were not included, due to their inherent interdependence. All non-correlated variables were included in the initial model and, for each set of correlated variables (vessel-related and MMO-related), we compared candidate models using the maximum likelihood (ML) and the Akaike Information Criterion (AIC; Zuur, et al., 2009; Pedersen et al., 2019). The final model was then refitted using the restricted maximum likelihood (REML) method (Zuur et al., 2009; Pedersen et al., 2019). All smooth terms were fitted with 4 splines (k = 4; Qian, 2016). Lastly, the best-fitting model was summarized using the function “summary(gam)”, and diagnostic plots were performed with the “gam.check” function. Influential data points and the correlation between the model residuals and explanatory variables were also checked. Then, to assess the model’s goodness of fit, a scatter plot of observed versus predicted values was used (Piñeiro et al., 2008). Concurvity values were used to assess potential non-linear dependencies among smooth terms (Amodio et al., 2014). All GAM analyses were developed with the “mgcv” R package.

## Results

### Descriptive and spatial analysis

A total of 1226 whale sightings were recorded, with 12 species and one genus-level identified (Table S1). CEs occurred across the entire study area, with a higher occurrence of NMEs between Madeira and mainland Portugal (Fig. 1A). A higher proportion of CEs in relation to the number of sightings was observed in the same region (Fig. 1B).

Of all whale sightings, 483 (39.4%) were classified as CEs, with 459 as SEs (37.4% of all sightings and 95.0% of all CEs) and 24 as NMEs (2.0% and 5.0%, respectively). The species most frequently involved in CEs were *Physeter macrocephalus* (n = 67, corresponding to 13.9% of all CEs), *Ziphius cavirostris* (n = 57, 11.8%), and *Balaenoptera acutorostrata* (n = 42, 8.7%), which were also the species most frequently sighted. Among these, *Z. cavirostris* showed the highest proportion of CEs relative to its total number of sightings (61.3%; Fig. 2A and Table S1).

**Figure 2.**
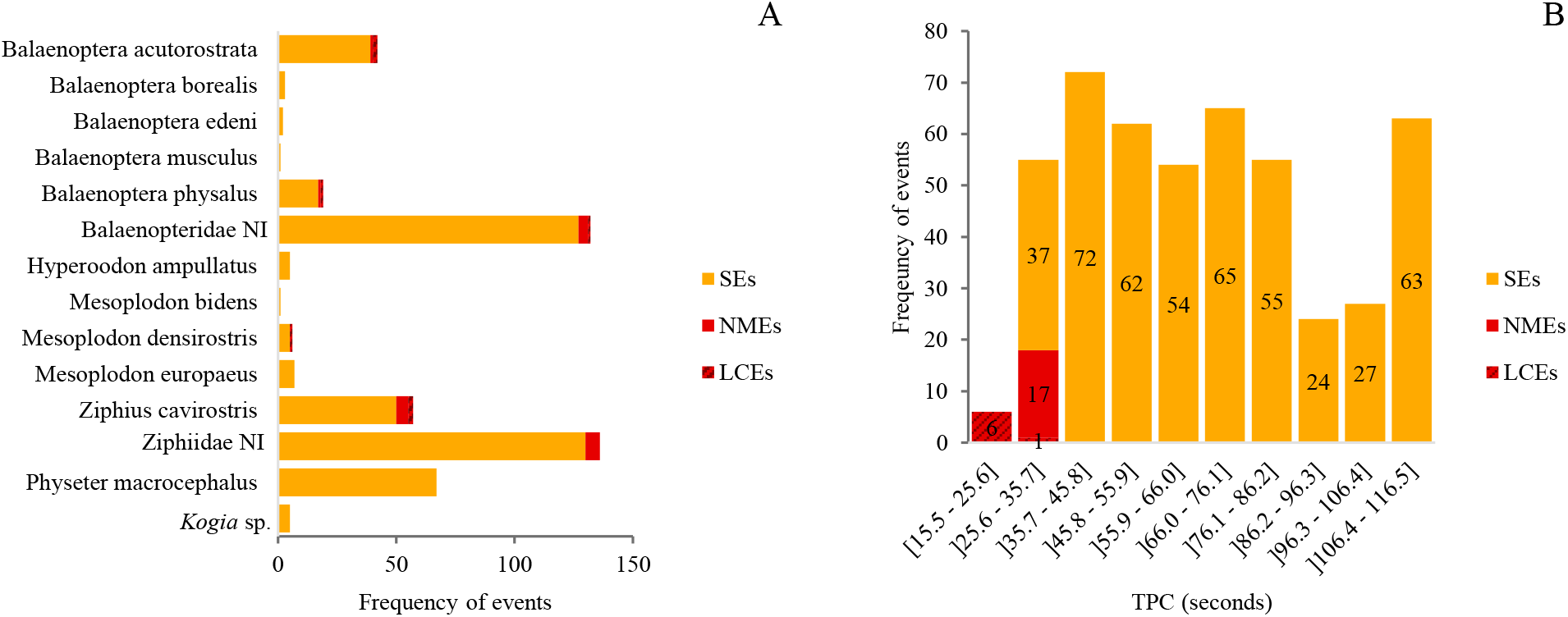
Histograms showing (A) the taxa involved in Surprise Encounters (SEs) and Near-Miss Events (NMEs); and (B) the distribution of Time to Potential Collision (TPC) of SEs and NMEs, collectively referred to as Close Encounters (CEs). Data were collected during cetacean monitoring surveys conducted between 2012 and 2024 in the Eastern North Atlantic, using cargo ships and oceanographic vessels as research platforms.

SEs involved 11 out of the 12 species sighted, whereas NMEs were associated with only four species: *B. acutorostrata, Balaenoptera physalus, Mesoplodon densirostris* and *Z. cavirostris* (Fig. 2A). Ziphiidae was the family most frequently associated with CEs, comprising a total of 198 SEs (43.1% of all SEs) and 14 NMEs (58.3% of all NMEs). Although not involved in any NME, *P. macrocephalus* was the species most involved with SEs (14.6% of all SEs), followed by *Z. cavirostris* (10.9%) and *B. acutorostrata* (8.5%). The *Z. cavirostris* accounted for the majority of NMEs (29.2% of all NMEs). Seven NMEs were classified as LCEs, and, in three of them, individuals exhibited an indifferent reaction towards the vessel (Fig. 2A and Table S1).

The TPC of CEs ranged from 15.5 to 116.5s, with a median of 63.3s. The most frequent TPC interval was 37.7 – 45.8s, accounting for 72 CEs (14.9% of CEs), and only six CEs (1.2%) had a TPC below 25.6s (Fig. 2B). Moreover, the shortest TPC was 15.5s, recorded for a *B. acutorostrata* sighting, closely followed by a *M. densirostris* sighting with a TPC of 15.8s.

Most whales involved in CEs were lone individuals (59.42% of all CEs), as were the majority involved in SEs (58.8% of all SEs) and NMEs (70.8% of all NMEs; Table S2). Furthermore, most of the individuals associated with a CE exhibited indifference towards the vessel (74.7% of all CEs), whilst 7.5% of CEs showed avoidance and 6.8% an approach. As for the NMEs, there were nine sightings in which the animal(s) showed avoidance (37.5% of all NMEs), eight that exhibited indifference (33.3%), and three that presented an approaching response (12.5%; Table S3).

### Modeling TPC

As there was a statistically significant difference between the TPC during on-effort and off-effort sightings (p-value = 0.01479), only the TPC of 885 on-effort sightings were included in the model. GVIF values of the explanatory variables were all below the established threshold, so all the variables were kept (Table S4). The explanatory variables retained in the final best model included sea state, wind state, visibility, taxonomic family, group size, vessel, LEO evaluation, and MEO evaluation, explaining 32% of the variation in TPC (Table 1).

**Table 1.** Results from the best-fitted Generalized Additive Model (GAM) developed to evaluate the influence of detectability-related variables on Time to Potential Collision (TPC) of on-effort whale sightings recorded between 2012 and 2024 in the Eastern North Atlantic, using cargo ships and oceanographic vessels as research platforms.

| n = 885; Deviance explained = 31.6%; R-Square (adjusted)= 0.231; -REML=5317.8; Scale est=0.618 |  |  |  |  |
| --- | --- | --- | --- | --- |
| Variable | EDF | Reference DF | F | P-value |
| s(group_size) | 0.6663 | 3 | 0.326 | 0.22052 |
| s(leo_evaluation) | 1.2727 | 3 | 1.001 | 0.09156 |
| s(meo_evaluation) | 1.6238 | 3 | 3.037 | 0.00314 |
| Variable | DF | F | P-value |  |
| sea_state | 3 | 1.023 | 0.3815 |  |
| wind_state | 3 | 0.741 | 0.5278 |  |
| visibility | 2 | 3.791 | 0.0229 |  |
| taxonomic_family | 2 | 80.426 | <2e-16 |  |
| vessel | 4 | 19.595 | 1.88e-15 |  |

When analyzing the fitted functions (Fig. 3), TPC generally decreased until a sea state of 2, after which it started to increase. The same trend was observed with wind state, whereas TPC increased with better visibility. TPC associated with Balaenopteridae and Physeteridae sightings was similar, while the Ziphiidae sightings presented a much lower TPC. TPC slightly increased with group size, up to five individuals, with an increasingly wide confidence interval thereafter, which limits the ability to draw robust conclusions. Regarding the vessel, TPC was higher in the DC vessel and generally lower in cargo ships, with IN holding the lowest TPC. Lastly, when considering the MMOs experience, TPC generally increased with higher LEO evaluations, and more markedly with higher MEO evaluations. Overall, significantly lower TPC values were observed with poor visibility conditions, beaked whales, cargo ships, and less experienced MMOs.

**Figure 3.**
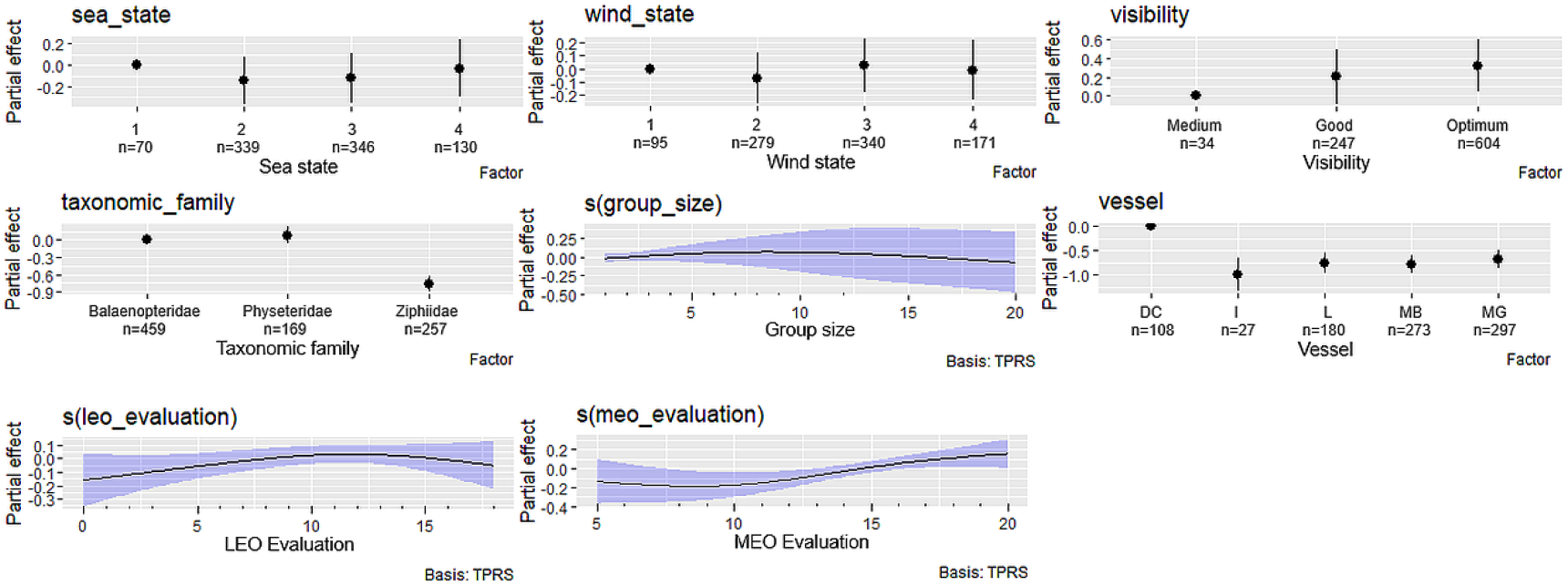
Plots of the final Generalized Additive Model (GAM) developed to evaluate the influence of detectability-related variables on Time to Potential Collision (TPC) of on-effort whale sightings recorded between 2012 and 2024 in the Eastern North Atlantic, using cargo ships and oceanographic vessels as research platforms. IN – Insular. DC – NRP Almirante D. Carlos I. GC – NRP Almirante Gago Coutinho. LA – Lagoa. MB – Monte Brasil. MG – Monte da Guia. LEO evaluation – Evaluation of the Least Experienced Observer on board. MEO evaluation – Evaluation of the Most Experienced Observer on board. Sea state – sea state classified according to the Douglas scale. Visibility – visibility states classified as medium (1 – 4 km), good (4 – 10 km), optimum (>10 km). Wind State - wind state classified according to the Beaufort scale.

Diagnostic plots, evaluation of model performance and Goodness of Fit, and concurvity values produced satisfactory results (Figs. S3-S6 and Tables S5-6).

## Discussion

### Defining Close Encounters

In this study, a speed-sensitive approach to estimate CEs is proposed, based on TPC rather than a fixed distance threshold. Including the influence of vessel speed on the definition of CEs is essential, as faster traveling speeds increase the likelihood of vessel-whale strikes (Vanderlaan & Taggart, 2007; Conn & Silber, 2013; Martin et al., 2016). Thus, when relying solely on distance to classify CEs, the actual risk of the whale being struck by the vessel is not being entirely considered, as it varies significantly with speed, and so does the time available for either the vessel or the whale to take evasive action. Setting thresholds based on TPC allows a more dynamic and risk-based classification of CEs, especially when considering a diverse array of speeds, as is the case of this study. Furthermore, these speed-dependent thresholds were discussed by Stack et al. (2016) and represent critical measurements of the time available to react and avoid collision. This methodology can further improve consistency when comparing CEs across vessels and studies, by considering each vessel’s traveling speed.

Nonetheless, using TPC as a metric has limitations. For instance, the difficulty in determining TPC *in situ* and, therefore, identifying a CE in real time. It is therefore advisable to record, in the field, the measurements required to calculate TPC, and subsequent CEs’ identification, specifically Vessel-to-Whale Distance and vessel speed. Then, for timely action at sea, a good strategy is to consider the vessels’ average cruise speed and estimate a distance range to the whale (multiplying vessel speed by the TPC threshold), upon which precautionary measures must be taken. The determination of this distance range should account for reaction time and vessel’s maneuverability and consider if it is sufficient to re-route (and/or reduce speed) and avoid a potential collision. For example, the TPC associated with an SE is of ∼2 minutes, which is a very low reaction time, likely insufficient to take evasive measures in most cases. Specifically in this study, the median TPC of CEs was approximately one minute, which may be insufficient time for the crew to react and does not allow much time for large ships (i.e., cargo ships) to maneuver. Thus, although the TPC thresholds here defined can be used analytically to characterize collision risk, they are not intended as operational guidance for ship handling. Real-time responses at sea are constrained by vessel maneuverability, traffic and meteorological conditions, and crew situational awareness, and must comply with applicable navigation regulations and established safety procedures.

Upon an imminent strike, implementing precautionary measures can be challenging, as decisions to avoid a whale often must be made within seconds (Ritter & Panigada, 2019). Moreover, predicting a whale’s trajectory and response towards an approaching vessel remains challenging, which limits the ability to avoid all potential strikes (Ritter, 2010; Wiley et al., 2016; Ritter & Panigada, 2019; Dunlop, 2024; Blondin et al, 2025). There are documented cases of vessel-whale NMEs even when the animal was seen well in advance (Ritter, 2010). This might suggest that even if there is enough time to maneuver the vessel away from the whale, there are still challenges. In fact, a linear vessel’s course, constant vessel speed, and a fixed whale position are often assumed, but real conditions (e.g., sea state fluctuations, movement/animal response) introduce some variability in the Vessel-to-Whale distance and consequently lead to a certain level of uncertainty in TPC. As such, the definition of a conservative TPC threshold upon which to act is an essential approach. If used as a basis to take evasive actions, TPC thresholds can help alert the crew to take timely precautionary measures and effectively avoid collisions. In summary, TPC-based mitigation measures (e.g., re-route and/or reduce speed) should be adapted to the situation and scaled to the level of risk, accounting for: distance of the animals, location of the sighting (bow or abeam), vessel maneuverability, and response of the animals (indifferent or aware).

### Descriptive and Spatial Analysis

A high percentage of sightings were classified as CEs, as the taken definition was quite conservative, ensuring that all possible CEs were included. Despite the broadness of this definition, any whale near a vessel can potentially be involved in any type of vessel-whale interaction (CEs and/or collision), under certain and specific conditions (Stack et al., 2016). In this study, only 2.0% of all sightings were classified as a NME, involving 4 identified species. This low percentage is comparable to findings reported by other authors, despite the different methodologies and thresholds (Richardson et al., 2011; Stack et al., 2013; David et al., 2022). Even though NMEs are considered somewhat rare, these are cases of very high collision risk. Among the identified NMEs, seven were classified as LCEs. Although none of them was directly confirmed as strikes, one can never be certain, as direct observations of collisions are inherently difficult from this type of vessel, and these events are the highest-risk scenario. Moreover, even if no whale suffered a strike, LCEs are expected to, at least, induce high levels of stress (Laist et al., 2001; David et al., 2022).

This study is limited to daylight hours, so it is not possible to account for the sightings and CEs that occur at nighttime, and it is very unlikely that any whale at risk is detected and avoided by the crew during these low visibility periods. This highlights even further the probable underestimation of CEs and collisions (Calambokidis et al., 2019; David et al., 2022). Moreover, a large proportion of non-identified (NI) whales (∼55.5%) were involved in CEs, suggesting that available data underestimates how often certain species are affected. Previous analyses have identified *B. physalus, Megaptera novaeangliae, B. edeni, P. macrocephalus*, and *B. acutorostrata* among the species most frequently involved in ship strikes (Laist et al., 2001; Winkler et al., 2020). Within this work, *P. macrocephalus, Z. cavirostris*, and *B. acutorostrata* were the species most associated with CEs. Considering the correlation between CEs and collision risk, the high occurrence of CEs associated with *P. macrocephalus* and *B. acutorostrata* is therefore expected. Specifically, ship-strike mortality of *P. macrocephalus* is particularly high in the Canary Islands (Carrillo & Ritter, 2010). On the other hand, *B. acutorostrata* is known to show much curiosity towards vessels and can sometimes appear suddenly at the bow (Perrin et al., 2018). This approaching response can lead to alarmed reactions by either the whale or crew members, imposing a greater risk of collision. Although most studies focus on great whales, beaked whales are also heavily affected by ship strikes (Schoeman et al., 2020; Winkler et al., 2020; Feyrer et al., 2024). In fact, most CEs involved Ziphiidae, suggesting a high risk of collision for these whales. As these animals inhabit extremely deep oceanic waters, there may exist a large overlap with the routes in this study, as most effort was in deep offshore areas (Mead, 2009). Beaked whales are often abundant near some seamounts and canyons, which are common features in the ENA, and this can also potentially contribute to the overlap (MacLeod & D’Amico, 2006; Mason, 2009; Moors-Murphy, 2014). Moreover, despite not spending much time at the surface, it is possible that their need for several breaths after long dives induces some level of anaerobic physiological stress, potentially diminishing their avoidance skills and putting them at a great collision risk (Van Waerebeek et al., 2007).

Previous studies suggest that a high proportion of CEs is often associated with lone individuals, rather than groups of two or more whales, which is consistent with the findings of the present study (Stack et al., 2013; Currie et al., 2017). For instance, Ritter (2010) described an NME involving two whales, in which the second whale only apparently noticed the approaching vessel when the first whale got startled. This alarmed reaction can suggest that lone whales may not always be aware of passing vessels and can benefit from social awareness in detecting potential threats (Ritter, 2010).

The high percentage of individuals associated with CEs, and specifically NMEs, who seemed indifferent to the vessel’s passage aligns with the literature. Whales who display an apparent indifferent response are more likely to be approached by a vessel, therefore being involved in a CE, and potentially being hit (Dolman et al., 2006; Ritter et al., 2016; Ritter & Panigada, 2019). Behaviors such as feeding, socializing, or resting are often associated with some indifference to the surroundings by the individuals, thus posing a higher risk for the animals (Dolman et al., 2006; Ritter et al., 2016; Ritter & Panigada, 2019; Blondin et al., 2025). On the other hand, the nine recorded NMEs in which whales exhibited an avoiding response towards the vessel might suggest that the ship might have been close enough for the whales to be aware of its approach and provoked an alarming and avoiding reaction from the individuals. This reaction, although potentially beneficial to avoid a collision, may indicate an increase in stress levels.

### Modeling TPC in relation to detectability factors

Although sea and wind states were included in the best explanatory model, they were not significant and their trend was less evident. Nevertheless, other studies have already pointed out that it can be challenging to spot and identify cetaceans under adverse meteorological conditions (Evans & Hammond, 2004; Richardson et al., 2011; Stack et al., 2016; Williams et al., 2016; Oliveira-Rodrigues et al., 2022). Even when considering only on-effort sightings, collected under favorable weather conditions, lower visibility still negatively influenced TPC. This leads to a reduced capacity to detect whales, and consequently a higher risk of a CE occurring.

Beaked whales are some of the most elusive cetacean species, making them difficult to detect, potentially increasing the risk of a collision (Mead, 2009; Ballance, 2018). This pattern is consistent with the higher frequency of these species associated with CEs in the present study, and with the significantly lower TPC associated with this taxa. Moreover, although this trend was not clearly evident in our results due to low sample size for larger groups, it is typically more challenging to detect smaller groups, especially when considering elusive species (Evans & Hammond, 2004; Williams et al., 2016; Oliveira-Rodrigues et al., 2022). In these situations, the MMOs may only sight the whales when they are already close to the vessel, leaving less time to avoid them.

Overall, TPC was higher in oceanographic vessels compared to cargo ships. Within the context of this study, these variations are possibly associated with the overall larger size and high traveling speeds associated with cargo ships. At faster speeds, the whale is detected closer to the vessel, causing a lower TPC and allowing less time to implement avoidance maneuvers (Vanderlaan & Taggart, 2007; Stack et al., 2013; Martin et al., 2016; Keen et al., 2022; Blondin et al., 2025). In fact, the recorded NMEs occurred exclusively with cargo ships. Despite this apparent trend between vessel speed and TPC, other vessels’ characteristics (e.g., overall length, platform height, distance to the bow, and tonnage) may have an influence, as they can affect detectability and sighting rates, and inherently collision risk (Evans & Hammond, 2004; Ritter et al., 2016). In fact, when the observer spots the animal in cargo ships, the Vessel-to-Whale Distance is already much lower due to the greater vessel’s length and distance to the bow, as the observer is positioned on the stern of the vessel. Moreover, among the cargo ships, IN was associated with the lowest TPC, followed by LA and MB, and lastly MG. Despite traveling at similar speeds, IN and LA have a lower observation deck, which can decrease detectability. The view towards the bow is often obstructed by containers, which may rise above the observation deck and limit visibility.

Lastly, an experienced MMO is likely more capable of spotting and detecting cetaceans at farther distances, thus influencing TPC. In this study, a higher TPC was associated with more experienced MMOs, reinforcing their importance on board. Employment of MMOs onboard consists of a real-time effective mitigation measure, as they can improve detection rates and support mitigation of vessel-whale collisions, by alerting the crew while increasing their awareness on this issue (potentially also increasing their willingness to take on evasive maneuvers; Carrillo & Ritter, 2010; Weinrich et al., 2010; Todd et al., 2015; Flynn & Calambokidis, 2019). As such, the employment of trained and dedicated MMOs should be established in mandatory guidelines, especially where collision risk is exceedingly high, and on board large vessels that travel at high speeds. However, MMOs’ efficiency depends on daylight and decreases with fatigue and, in the presence of poor weather conditions, detecting cetaceans becomes even more challenging (Harwood & Joynt, 2009; Oliveira-Rodrigues et al., 2022). As such, automatic detection systems, such as thermal infrared imaging devices, are being employed. Although useful, these methodologies are costly and still present some limitations, such as the generation of large datasets for calibration, malfunctions, occasional inaccuracies, and challenges associated with automatic detections and alert systems (Horton et al., 2017; Baille & Zitterbart, 2022). Whether these methodologies can function as an effective mitigation measure depends on certain parameters, such as vessel speed and maneuverability, and the accuracy of the detection system (Baille & Zitterbart, 2022). The optimization and validation of these methods, as well as the correction of errors and malfunctions associated with them (and the need for their lower cost and technological accessibility), continues to require the presence of MMOs on board, hence their importance (Baille & Zitterbart, 2022). Despite some restraints, automatic detection systems can serve as good complementary techniques to MMOs and can potentially increase the detection of risk events, as well as TPC (IWC, 2011; Verfuss et al., 2016; Horton et al., 2017; Zitterbart et al., 2020; Baille & Zitterbart, 2022). Therefore, further efforts should be made to integrate both methods.

Our results confirm the widespread occurrence of CEs within the ENA, highlighting a potential risk of collisions. The thresholds defined here to characterize CEs, based on the time available to react, represent a relevant effort to standardize CEs’ quantification, which remains challenging. Using TPC as a metric allows CEs to be recorded under a risk-based classification, which enables their quantification to be compared across different studies. We therefore suggest future studies to adopt these time-based thresholds, considering the role of vessel speed in the likelihood of a collision. The findings also demonstrate that whale characteristics influence the likelihood of CEs, with beaked whales being the most affected taxa, underscoring the need to study collision risk on both medium- and large-sized whales. Furthermore, collision risk is influenced by vessel characteristics and meteorological conditions. Variables that negatively influence detectability led to lower TPC values, thereby increasing the risk to whales. These results acknowledge the importance of measures that enhance detectability, such as the employment of MMOs. Furthermore, initiatives to raise awareness of this anthropogenic pressure should be the first step towards collision mitigation. Alerting the crew for this issue and co-create preventive measures is of extreme relevance to foster collaboration and willingness to undertake appropriate, timely actions upon collision risk. When MMOs are unavailable, a trained crew could aid in data collection, identification of risk situations, and collision prevention. To achieve this, close cooperation and effective communication among researchers, policymakers, and the maritime community need to be implemented. In fact, the best conservation and management strategies should result from a concerted effort from all sectors.

## Acknowledgments

The data were collected within the CIIMAR long-term cetacean monitoring programme of the CETUS project. We thank Grupo ETE – Transinsular (Portuguese shipping company) and the Hydrographic Institute of the Portuguese Navy, for their logistical support during the data collection, through their established collaborations with CIIMAR within the CETUS project. We are also grateful to all vessel crews for their support in accommodating observers on board. Lastly, we thank all the volunteers who contributed to the project during monitoring campaigns as dedicated MMOs. They are Adriana Melo, Ágatha Gil, Alberto Río-Pérez, Alexandra Pires, Alexandre Branco, Amber Coleman, Amélie Gadbois, Ana Ayres, Andreia Pereira, Anicée Lombal, Anja Badenas, António Nogueira, Anxo Gende, Ayca Eleman, Bárbara Matos, Beatriz Filipe, Carmen Escobar, Catarina Fonseca, Catarina Morgado, Cláudia Ferreira, Cláudia Rodrigues, Corina Peter, Diana Fonseca, Eduardo Lima, Esteban Iglesias, Fadia Al Abbar, Firat Hayta, Francisco Cardeal, Iolanda Silva, Jack Ball, Joana Romero, Joana Silva, Jorge Silva, Judit Miquel, Julia Heiler, Julia Müller, Juliana Moron, Julien Freyer, Lara Meraz, Laura García, Layla Mpinou, Leonardo Berninsone, Lidia Lado, Luís Afonso, Mafalda Correia, Margot Paris, Maria Ballesteros, María Jiménez, Marieta Mihova, Mayara Gomes, Mieke Weyn, Nádia Silva, Natalia Jimenez, Nicolas Blanc, Oleh Danduriants, Olga Azevedo, Patrícia Carvalho, Paula Carreño, Paula Silva, Pedro Fernandes, Pedro Silva, Pedro Xavier, Philippa Wright, Raúl Valente, Rebeca Velasco, Rhiannon Nichol, Rita Figueiras, Rita Santos, Roosevelt Gutiérrez, Ryan Matthews, Sandra Más, Santiago Otero, Sara Araújo, Sara Pastorino, Sarah Drake, Sofia Silva, Suzana Martins, Svenja Halfter, Tanja Schwanck, Tara Callahan, Tiago Brandão, Torcuato Mantas, Valéria Aragão, Vasco Gomes, Verónica Belchior, Victoria Hope, and Zara Valquíria.

