## Supplemental Figure S1 for "Time to Potential Collision: A Dynamic Approach To Study Vessel-Whale Close Encounters"

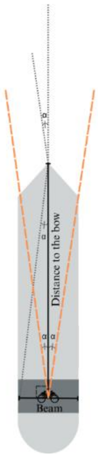


**Figure S1.** Schematic representation of the method used to determine the angular range corresponding to the vessel's bow area (α).
