## Supplemental Figure S2 for "Time to Potential Collision: A Dynamic Approach To Study Vessel-Whale Close Encounters"

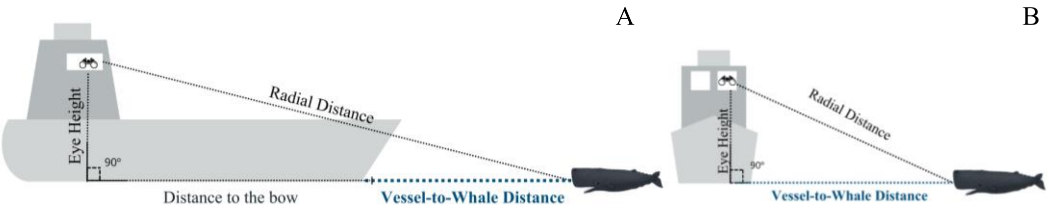


**Figure S2.** Schematic representation of the method used to calculate the Vessel-to-Whale Distance for sightings occurring at the vessel's bow (defined as between -6º and 6º for cargo ships and between -23º and 23º for oceanographic vessels; A) and outside the vessel's bow angular range (i.e., sightings abeam; B).
