## Supplemental Figure S3 for "Time to Potential Collision: A Dynamic Approach To Study Vessel-Whale Close Encounters"

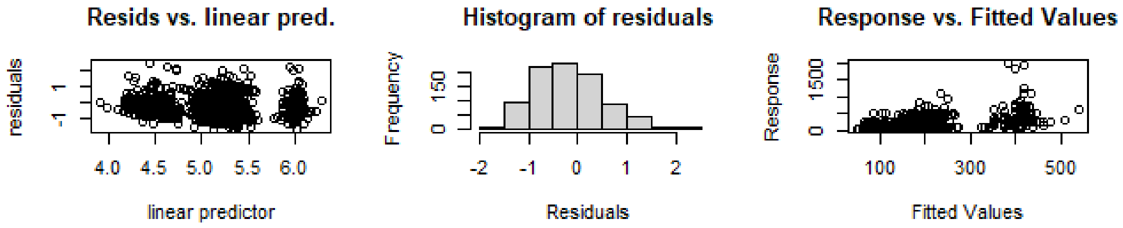


**Figure S3.** Generalized Additive Model (GAM) Check Plots of the model developed to evaluate the influence of detectability-related variables on Time to Potential Collision (TPC) of on-effort whale sightings recorded between 2012 and 2024 in the Eastern North Atlantic, using cargo ships and oceanographic vessels as research platforms.
