## Supplemental Figure S5 for "Time to Potential Collision: A Dynamic Approach To Study Vessel-Whale Close Encounters"

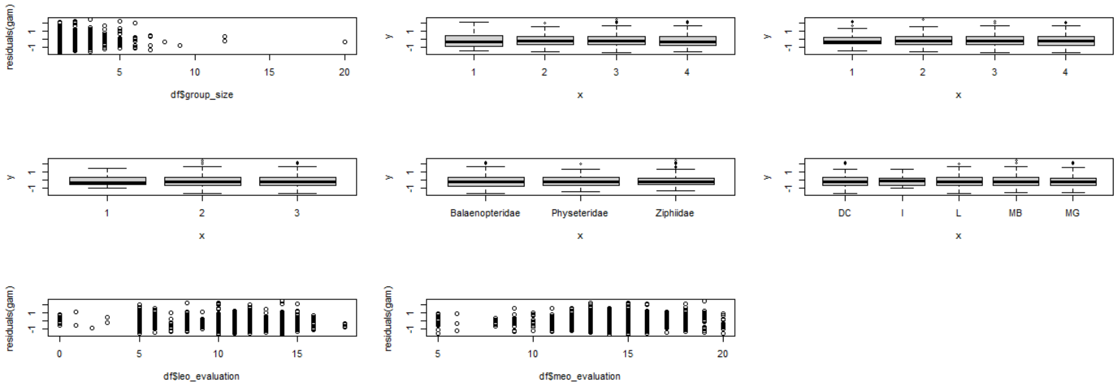


**Figure S5.** Residuals plots of the explanatory variables included in the model developed to evaluate the influence of detectability-related variables on Time to Potential Collision (TPC) of on-effort whale sightings recorded between 2012 and 2024 in the Eastern North Atlantic, using cargo ships and oceanographic vessels as research platforms. IN – Insular. DC – NRP Almirante D. Carlos I. GC – NRP Almirante Gago Coutinho. LA – Lagoa. MB – Monte Brasil. MG – Monte da Guia.
