## Supplemental Figure S6 for "Time to Potential Collision: A Dynamic Approach To Study Vessel-Whale Close Encounters"

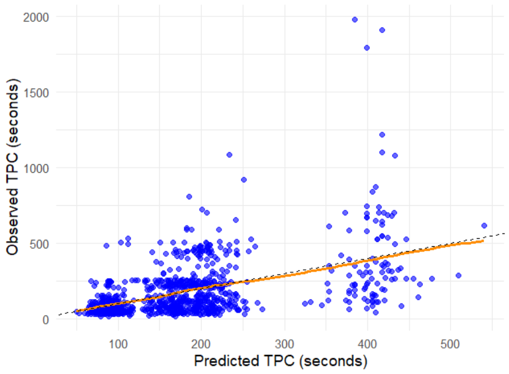


**Figure S6.** Goodness of Fit of the Generalized Additive Model (GAM) developed to evaluate the influence of detectability-related variables on Time to Potential Collision (TPC) of on-effort whale sightings recorded between 2012 and 2024 in the Eastern North Atlantic, using cargo ships and oceanographic vessels as research platforms.
