## Supplemental Table S1 for "Time to Potential Collision: A Dynamic Approach To Study Vessel-Whale Close Encounters"

**Table S1.** Summary of whale sightings and Close Encounters (CEs), including Surprise Encounters (SEs), Near-Miss Events (NMEs), and Likely-Collision Events (LCEs), recorded during cetacean monitoring surveys conducted between 2012 and 2024 in the Eastern North Atlantic, using cargo ships and oceanographic vessels as research platforms.

| Taxa | Whale sightings | | CEs | | | SEs | | NMEs | | |
|---|---|---|---|---|---|---|---|---|---|---|
| | Frequency | % (in relation to all whale sightings) | Frequency | % (in relation to the total of CEs) | % (in relation to the sightings within the taxa) | Frequency | % (in relation to the total of SEs) | Frequency | % (in relation to the total of NMEs) | LCEs |
| *Balaenoptera acutorostrata* | 97 | 7.9 | 42 | 8.7 | 43.3 | 39 | 8.5 | 3 | 12.5 | 2 |
| *Balaenoptera borealis* | 8 | 0.7 | 3 | 0.6 | 37.5 | 3 | 0.7 | 0 | 0.0 | |
| *Balaenoptera edeni* | 6 | 0.5 | 2 | 0.4 | 33.3 | 2 | 0.4 | 0 | 0.0 | |
| *Balaenoptera musculus* | 3 | 0.2 | 1 | 0.2 | 33.3 | 1 | 0.2 | 0 | 0.0 | |
| *Balaenoptera physalus* | 57 | 4.6 | 19 | 3.9 | 33.3 | 17 | 3.7 | 2 | 8.3 | 1 |
| *Megaptera novaeangliae* | 9 | 0.7 | 0 | 0.0 | 0.0 | 0 | 0.0 | 0 | 0.0 | |
| Balaenopteridae NI | 474 | 38.7 | 132 | 27.3 | 27.8 | 127 | 27.7 | 5 | 20.8 | 1 |
| Balaenopteridae | 654 | 53.3 | 199 | 41.2 | 30.4 | 189 | 41.2 | 10 | 41.7 | 4 |
| *Hyperoodon ampullatus* | 7 | 0.6 | 5 | 1.0 | 71.4 | 5 | 1.1 | 0 | 0.0 | |
| *Mesoplodon bidens* | 3 | 0.2 | 1 | 0.2 | 33.3 | 1 | 0.2 | 0 | 0.0 | |
| *Mesoplodon densirostris* | 8 | 0.7 | 6 | 1.2 | 75.0 | 5 | 1.1 | 1 | 4.2 | 1 |
| *Mesoplodon europaeus* | 9 | 0.7 | 7 | 1.4 | 77.8 | 7 | 1.5 | 0 | 0.0 | |

| | | | | | | | | | |
|---|---|---|---|---|---|---|---|---|---|
| *Ziphius cavirostris* | 93 | 7.6 | 57 | 11.8 | 61.3 | 50 | 10.9 | 7 | 29.2 | 2 |
| Ziphiidae NI | 217 | 17.7 | 136 | 28.2 | 62.7 | 130 | 28.3 | 6 | 25.0 | |
| Ziphiidae | 337 | 27.5 | 212 | 43.9 | 62.9 | 198 | 43.1 | 14 | 58.3 | 3 |
| *Physeter macrocephalus* | 227 | 18.5 | 67 | 13.9 | 29.5 | 67 | 14.6 | 0 | 0.0 | |
| Physeteridae | 227 | 18.5 | 67 | 13.9 | 29.5 | 67 | 14.6 | 0 | 0.0 | |
| *Kogia* sp. | 8 | 0.7 | 5 | 1.0 | 62.5 | 5 | 1.1 | 0 | 0.0 | |
| Kogiidae | 8 | 0.7 | 5 | 1.0 | 62.5 | 5 | 1.1 | 0 | 0.0 | |
| Total | 1226 | | 483 | | | 459 | | 24 | | 7 |
