## Supplemental Table S2 for "Time to Potential Collision: A Dynamic Approach To Study Vessel-Whale Close Encounters"

**Table S2.** Frequency of Close Encounters (CEs), including Surprise Encounters (SEs) and Near Miss Events (NMEs), according to the best estimate of the group size, of data collected during cetacean monitoring surveys conducted between 2012 and 2024 in the Eastern North Atlantic, using cargo ships and oceanographic vessels as research platforms. Percentages indicate the proportion of SEs, NMEs and total CEs relative to the total number of events in each category, respectively.

| Group size (best estimate) | CEs | | Total |
|---|---|---|---|
| | SEs | NMEs | |
| 1 | 270 (58.8%) | 17 (70.8%) | 287 (59.4%) |
| 2 | 107 (23.3%) | 4 (16.7%) | 111 (23.0%) |
| 3 | 46 (10.0%) | 1 (4.2%) | 47 (9.7%) |
| 4 | 16 (3.5%) | 1 (4.2%) | 17 (3.5%) |
| 5 | 11 (2.4%) | 0 | 11 (2.3%) |
| 6 | 3 (0.7%) | 1 (4.2%) | 4 (0.8%) |
| 7 | 3 (0.7%) | 0 | 3 (0.6%) |
| 8 | 1 (0.2%) | 0 | 1 (0.2%) |
| 9 | 1 (0.2%) | 0 | 1 (0.2%) |
| 12 | 1 (0.2%) | 0 | 1 (0.2%) |
