## Supplemental Table S4 for "Time to Potential Collision: A Dynamic Approach To Study Vessel-Whale Close Encounters"

**Table S4.** Generalized Variance Inflation Factor (GVIF) results of explanatory variables of the model developed to evaluate the influence of detectability-related variables on Time to Potential Collision (TPC) of on-effort whale sightings recorded between 2012 and 2024 in the Eastern North Atlantic, using cargo ships and oceanographic vessels as research platforms.

| Variable | GVIF |
|---|---|
| group_size | 1.124143 |
| sea_state | 1.687582 |
| wind_state | 1.647902 |
| visibility | 1.123159 |
| taxonomic_family | 1.191452 |
| leo_evaluation | 1.308582 |
| meo_evaluation | 1.339052 |
