## Supplemental Table S5 for "Time to Potential Collision: A Dynamic Approach To Study Vessel-Whale Close Encounters"

**Table S5.** Results of basis dimension (k) checking (with gam.check) of the Generalized Additive Model (GAM) developed to evaluate the influence of detectability-related variables on Time to Potential Collision (TPC) of on-effort whale sightings recorded between 2012 and 2024 in the Eastern North Atlantic, using cargo ships and oceanographic vessels as research platforms.

| Variable | K’ | EDF | K-index | P-value |
|---|---|---|---|---|
| s(group_size) | 3.00 | 0.666 | 0.92 | 0.25 |
| s(leo_evaluation) | 3.00 | 1.273 | 0.93 | 0.40 |
| s(meo_evaluation) | 3.00 | 1.624 | 0.91 | 0.17 |
