## Supplemental Table S6 for "Time to Potential Collision: A Dynamic Approach To Study Vessel-Whale Close Encounters"

**Table S6.** Concurvity values of the Generalized Additive Model (GAM) developed to evaluate the influence of detectability-related variables on Time to Potential Collision (TPC) of on-effort whale sightings recorded between 2012 and 2024 in the Eastern North Atlantic, using cargo ships and oceanographic vessels as research platforms.

| | s(group_size) | s(leo_evaluation) | s(meo_evaluation) |
|---|---|---|---|
| worst | 0.1525540 | 0.3061972 | 0.3710285 |
| observed | 0.1471336 | 0.1718969 | 0.3571375 |
| estimate | 0.1173315 | 0.2521247 | 0.3496867 |
